# Mechano-epigenetic regulation of extracellular matrix homeostasis via Yap and Taz

**DOI:** 10.1101/2022.07.11.499650

**Authors:** Dakota L. Jones, Ryan N. Daniels, Xi Jiang, Ryan C. Locke, Mary Kate Evans, Edward D. Bonnevie, Anjana Srikumar, Madhura P. Nijsure, Joel D. Boerckel, Robert L. Mauck, Nathaniel A. Dyment

## Abstract

Cells integrate mechanical cues to direct fate specification to maintain tissue function and homeostasis. While disruption of these cues is known to lead to aberrant cell behavior and chronic diseases, such as tendinopathies, the underlying mechanisms by which mechanical signals maintain cell function is not well understood. Here, we show using a novel model of tendon de-tensioning that loss of tensile cues *in vivo* acutely changes nuclear morphology, positioning, and expression of catabolic gene programs. Using paired ATAC/RNAseq, we further identify that a loss of cellular tension rapidly reduces chromatin accessibility in the vicinity of Yap/Taz genomic targets while also increasing expression of genes involved in matrix catabolism. Overexpression of Yap results in a reduction of chromatin accessibility at matrix catabolic gene loci, while also reducing transcriptional levels. Concordantly, depletion of Yap/Taz elevates matrix catabolic expression. Finally, we demonstrate that overexpression of Yap not only prevents the induction of a broad catabolic program following a loss of cellular tension, but also preserves the underlying chromatin state from force-induced alterations. Taken together, these results provide novel mechanistic details by which mechanical signals regulate tendon cell function to preserve matrix homeostasis through a Yap/Taz axis.

**Significance Statement:** Cells integrate mechanical signals to regulate biological outputs within tissues. These processes are required for tissue function and homeostasis. Here, we show how mechanical cues (e.g. tension) directs tendon cell function and fate at a transcriptional and epigenetic level. Furthermore, we show that disruption of these mechanical cues leads to a disease-like cell state, indicating these mechanosensitive pathways could be important for diseases driven by perturbed mechanical signaling, such as tendinopathy. Finally, we demonstrate that genetic perturbation of a single protein can preserve cell and chromatin state following a loss of tension, supporting novel avenues for the development of innovative mechano-therapeutics.

## Introduction

All tissues, and in particular those that are designed to bear load, sense and respond to physical forces. Indeed, mechanically-mediated tissue remodeling is a hallmark of musculoskeletal development, homeostasis, and disease progression (1). For example, tendons, a dense collagenous connective tissue whose function is to transmit tensile forces from muscle to bone, are subject to thousands of loading cycles each day as a consequence of muscle contraction during activities of daily living (2). Tendons maintain a state of residual tension on which these applied loads are superimposed (3). This “tensional homeostasis” is critical for maintaining tissue properties of tendons. Intrinsic tension arises from both direct cellular contractility and by residual stress in the tendon matrix. Both intrinsic and extrinsic tensile cues are thus central to tissue function, and loss of either can impact tissue homeostasis (4, 5). Loss of tensional homeostasis at the macroscale, as occurs with transection or periods of immobility, or at the microscale, as is the case when micro-damage occurs that locally interrupts for fibrous architecture of the tissue, alters load transmission to the endogenous fibroblast cell population. The loss of these tensile cues in turn perturbs the homeostatic function of tendon fibroblasts (5–7). Indeed, tendinopathic tissue show an imbalance in matrix turnover and at later stages tendon cells may actually differentiate along aberrant lineages such as cartilage and bone (8). While the degenerative end state of tendinopathy is well established, the early mechanistic cellular events that are mediated by changes in mechanical loading are not yet known.

Mechano-responsive signaling pathways are central regulators of fibroblast and other mesenchymal cells, controlling a spectrum of activities from proliferation, contractility, differentiation, and ECM remodeling (9–11). In tensile loading scenarios, mechanical cues are relayed through the contractile cytoskeleton to the nucleus to control transcriptional activities either indirectly through activation of other mechano-signaling pathways (e.g., stretch activated channels, mechano-responsive transcriptional co-activators such as YAP/TAZ) or directly (e.g. deformation of the nucleus, nuclear pores, and chromatin)(12–16). Whether through direct chromatin deformation or via mobilization of mechano-active factors, mechano-signaling pathways ultimately converge onto transcriptional regulators, which possess epigenetic and transcriptional activity to regulate cell function (17). *In situ* and *in vitro* studies show that disruption of mechanosignaling in tendon and other connective tissue fibroblasts alters their functionality (4, 5, 18, 19). Other work has shown that mechanical cues, such as matrix stiffness, can regulate chromatin accessibility and other epigenetic features in fibroblasts and other cell types, suggesting a direct role for these cues in regulating the epigenetic state and transcriptional receptiveness of these cells to mechano-activation (20–23). However, the transcriptional events that dictate how these cells respond to an acute loss of mechanical signals remains unknown.

To address this gap in knowledge, we evaluated tendon cell response to a loss of tensional homeostasis, *in vitro* and *in vivo*. Our data show that an acute loss of tension results in an increase in the expression of catabolic gene programs in tendon fibroblasts. Subsequent mechanistic epigenomic (ATACseq) and transcriptomic (RNAseq) analyses identified that cytoskeletal de-tensioning results in a global decrease in chromatin accessibility coincident with an increase in the expression of a matrix catabolic gene program. Unbiased analyses of ATACseq performed on de-tensioned tendon fibroblasts identified Yap/Taz/Tead targets as differentially regulated loci, suggesting that Yap/Taz/Tead may act as a transcriptional repressors of catabolic gene programs under homeostatic conditions. Depletion of Yap/Taz phenocopied the loss of cellular tension, instigating a broad catabolic program. Conversely, overexpression of Yap/Taz reduced chromatin accessibility and downstream expression of genes involved in matrix catabolism, even in the face of a loss of cellular tension. Not only did this prevent cells from initiating a catabolic program, overexpression of Yap/Taz preserved accessibility at target chromatin loci following loss of tension. Taken together, these data support that cellular tension, acting through Yap/Taz/Tead, promotes an anabolic state in tendon fibroblasts, and thus may serve as a potential therapeutic avenue to inhibit ECM loss in scenarios where macroscopic and microscopic tensional homeostasis is perturbed.

## Results

### Cellular tension negatively regulates extracellular matrix degradative pathways

To investigate the role of tensile cues in directing tendon-resident fibroblast function, we developed an *in vivo* loss-of-tension model in which we resected a distal segment of the flexor digitorum longus tendon (FDL) in the mouse forefoot, allowing the proximal portion of the tendon to be analyzed while minimizing effects from the early stages of healing (e.g., clot formation, granulation tissue) at the injury site (**Fig. 1a**). To examine the early response of the cell nucleus to the loss of tension, we quantified the nuclear aspect ratio and the dispersion (i.e., higher dispersion indicates cells/nuclei are more disorganized) of DAPI-stained tissue sections at 24 hours post-resection. Interestingly, acute loss of tension reduced nuclear aspect ratio and increased nuclear dispersion (**Fig. 1b, c**) in the resected tendons compared to uninjured contralateral controls. These data demonstrate that *in vivo* tensile cues are central to maintenance of overall nuclear shape and positioning.

**Figure 1:**
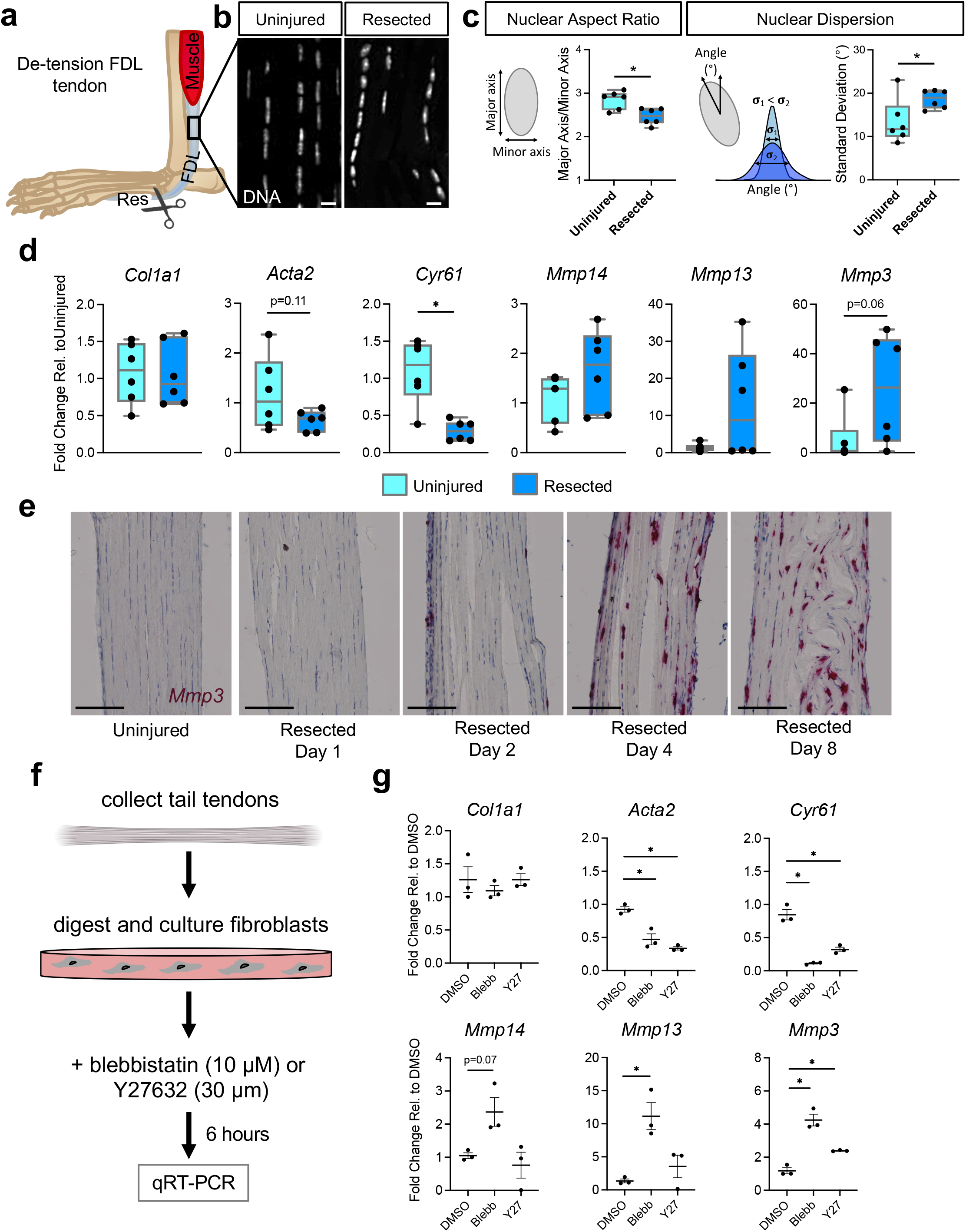
Pathologic acute mechano-responses of tendon fibroblasts to de-tensioning. (a) Schematic showing approach to de-tensioning the flexor digitorum longus (FDL) tendon *in vivo*. (b) DAPI staining depicting nuclear morphology following loss of tension (scale bar = 10μm). (c) Schematic and quantification of nuclear aspect ratio and nuclear dispersion angle of tendon fibroblasts 24 hours following loss of tension in the FDL (n=6 biological replicates). (d) qRT-PCR analysis of FDL tendons 24 hours following resection (n=6 biological replicates). (e) RNAScope analysis for Mmp3 over time following resection of the FDL tendon (scale bar = 100μm). (f) Schematic approach showing approach for isolation and culture/experimental conditions for tendon fibroblasts *in vitro*. (g) qRT-PCR analysis of tendon fibroblasts 6 hours following addition of blebbistatin (10 μM) and Y27632 (30 μM) (n=3 biological replicates). *P<0.05 evaluated by unpaired t-test. Error bars represent standard error.

To better understand how these early changes in nuclear shape might affect downstream gene expression, we performed qRT-PCR on tendons collected 24 hours after resection. While analysis of transcripts from the de-tensioned tendons showed no change in matrix anabolic genes (i.e., *Col1a1*), we observed an increase in the expression of matrix catabolism genes (i.e., matrix metalloproteinases (MMPs)), demonstrating that the acute early response to a loss of tensile cues is an upregulation of matrix degradative gene programs (**Fig. 1d)**. Interestingly, we also noted reduced expression of *Acta2* and *Cyr61*, established mechano-responsive genes (**Fig. 1d**), suggesting that tendon fibroblasts are responsive to the loss of tensile mechanical cues *in vivo*. Using RNAscope, we performed a time course experiment to better understand the kinetics of *Mmp3* expression following resection. Interestingly, we observed a progressive increase in *Mmp3* within the resident tendon cells over time (**Fig. 1e**) suggesting that the early acute response to a loss of tension leads to persistent changes in matrix catabolic gene expression. Collectively, these data demonstrate that tendon-resident fibroblasts are exquisitely sensitive to tensile cues within the tissue. Furthermore, de-tensioning results in acute changes in nuclear morphology, mechanical signaling, and increased expression of matrix catabolic genes, potentially leading to chronic matrix degradation.

To elucidate the short-term mechano-response which regulates this transcriptional program, we de-tensioned tendon fibroblasts *in vitro* using two small-molecule inhibitors of proteins involved in mechano-signaling, blebbistatin inhibiting myosin contraction and Y27632 inhibiting Rho kinase. Using freshly isolated tendon fibroblasts (**Fig. 1f**), we administered blebbistatin (10 μM) and separately Y27632 (30 μM) and analyzed a broad set of matrix catabolic genes by qRT-PCR. Similar to de-tensioning *in vivo*, we found marked upregulation of matrix catabolic genes, particularly following inhibition of myosin contractility but less so Rho kinase (**Fig. 1g**), demonstrating that cellular tension plays a key role in regulating expression of these pathways. Taken together, these data suggest that an early response to loss of tensile cues could be altered expression of matrix degradation pathways, a potentially critical step in the progression to chronic tendinopathy.

### Mechanical regulation of epigenomic and transcriptomic state of tendon fibroblasts

To investigate the transcriptional and epigenetic mechanisms through which intracellular tension mediates this extracellular matrix homeostasis, we administered blebbistatin (10 μM) for 6 hours and performed RNAseq and ATACseq. Following blebbistatin treatment, RNAseq analysis revealed increased transcripts from 570 genes and decreased transcripts from 462 genes (**Fig. 2a**). Gene ontology analysis of those gene lists identified that inhibition of cytoskeletal tension did indeed increase collagen catabolism pathways, confirming a role for cellular tension in regulating genes involved in extracellular matrix homeostasis (**Fig. 2b**).

**Figure 2:**
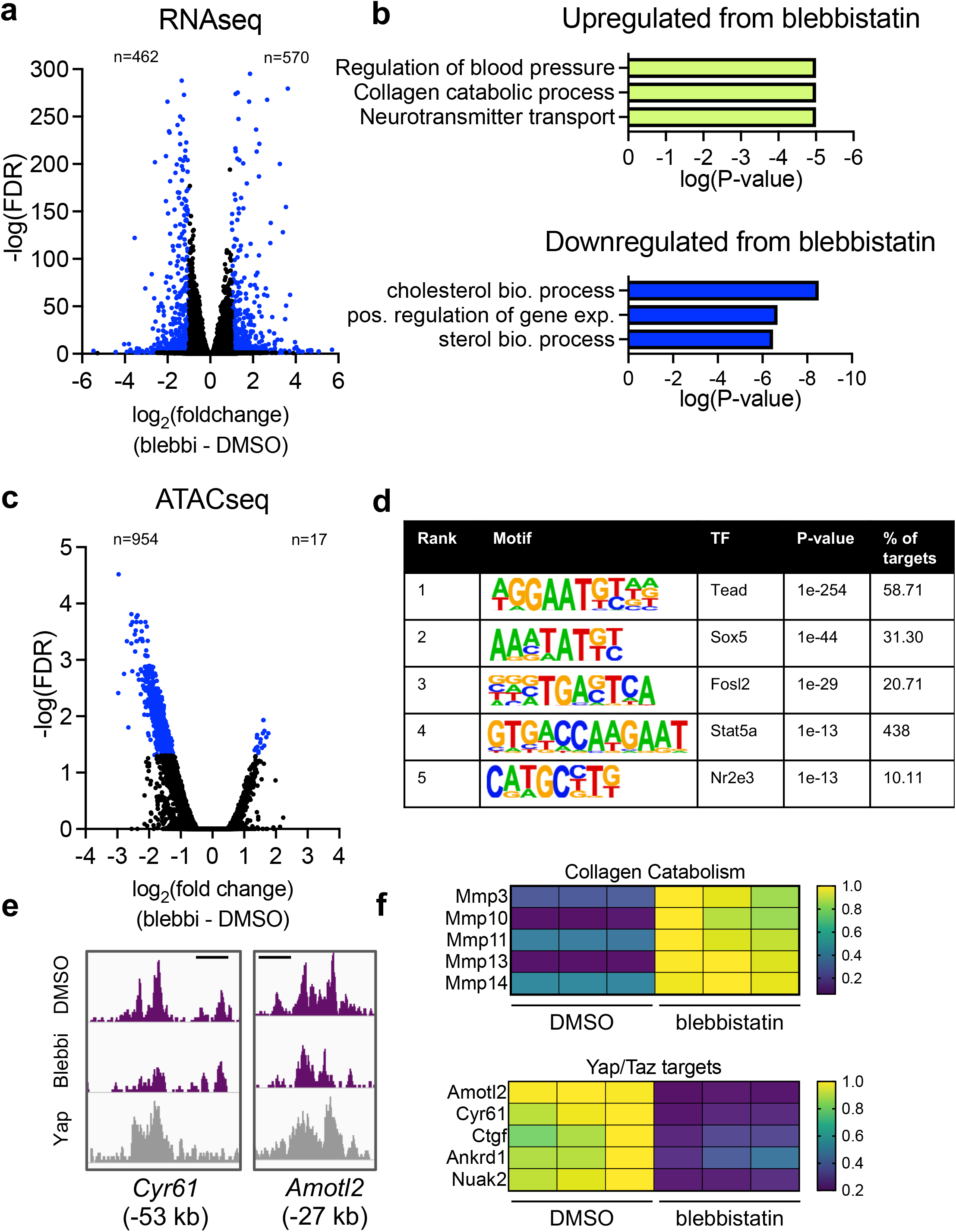
Cellular tension maintains epigenetic and transcriptional homeostasis in tendon fibroblasts. (a) Volcano plot from RNAseq analysis showing differentially expressed genes (adjusted p-value ≤ 0.05 and a fold-change of ≤ -1 or ≥ 1) in blue (n=3 biological replicates). (b) gene ontology analysis of gene sets identified in (a). (c) Volcano plot from ATACseq analysis showing differentially accessible genomic loci (adjusted p-value ≤ 0.05 and a fold-change of ≤ -1 or ≥ 1) in blue (n=3 biological replicates). (d) De novo transcription factor motif enrichment analysis from the genomic loci identified in (c). Transcription factors are ranked by statistical enrichment (p-value). (e) Representative changes in chromatin accessibility following blebbistatin treatment cross-referenced with publicly available Yap ChIPseq (GSE83863). Genomic distance represents distance from gene transcriptional start site (TSS). Scale bar represents 1kb. (f) Representative gene expression changes of collagen catabolism genes and Yap/Taz target genes following blebbistatin treatment.

Interestingly, ATACseq revealed that blebbistatin reduced accessibility at 954 chromatin loci and increased accessibility at only 17 chromatin loci. These results indicate that active cytoskeletal tension, on the whole, acts to positively regulate chromatin accessibility (**Fig. 2c**).

To identify potential transcriptional regulators driving these epigenetic and transcriptional changes, we performed *de novo* transcription factor motif enrichment analysis within the chromatin loci that were either increased or decreased in accessibility following blebbistatin treatment (**Fig. 2c**). This unbiased motif analysis identified Tead as the leading transcription factor (**Fig. 2d**). The Tead family of proteins function as transcriptional co-activators for Yap and Taz, which are major mechano-responsive transcription factors involved in cellular mechanosensing (12). This suggests that tensile cues could mediate extracellular matrix homeostasis through a Yap/Taz/Tead axis. Further supporting this idea, co-occupancy analysis of publicly available Yap ChIPseq data (GSE83863)(24) demonstrated a robust overlap of Yap occupancy with differentially accessible chromatin loci following blebbistatin treatment (**Fig. 2e**). Analysis of Yap/Taz target genes within our RNAseq data confirmed a reduction in transcripts of Yap/Taz target genes as well as an increase in transcripts of a broad group of MMPs and cathepsins (**Fig. 2f**, GSE207896). Collectively, these data demonstrate that loss of cellular tension disrupts chromatin state, modulating accessibility at Yap/Taz/Tead target loci, resulting in an increase in transcriptional activity of genes in matrix degradation pathways.

### Yap/Taz/Tead represses matrix degradation pathways

Given that the ATACseq analysis pointed to Yap and/or Taz as potential key mediators of the response to loss of cytoskeletal tension in tendon fibroblasts, we next used a B6.C-Tg(CMV-cre)1Cgn/J;TetO-Yap^S127A^; Gt(ROSA)26Sor^tm1(rtTA,EGFP)Nagy/J^ mouse (herein referred as YAP-CA) which overexpresses a mutated constitutively-active human YAP1 gene following doxycycline administration to further assess the role of Yap/Taz in these cells (25, 26). We isolated primary fibroblasts from tail tendons from adult YAP-CA mice and found robust YAP overexpression following 48 hours of doxycycline administration (10ng/mL) (**Fig. 3a**,**b**). To better understand the role of Yap/Taz in regulating chromatin state, we administered doxycycline for 48 hours and then performed ATACseq. ATACseq analysis revealed that overexpression of YAP resulted in an increase of accessibility at 7,297 chromatin loci and only decreased accessibility at 316 chromatin loci (**Fig. 3c**). These data are consistent with YAP maintaining chromatin “open-ness”, potentially acting as a pioneer factor directly regulating chromatin accessibility. As expected, *de novo* motif enrichment analysis within the chromatin loci that changed accessibility following YAP overexpression revealed Tead as the leading candidate, confirming a Yap/Taz/Tead axis as directly regulating chromatin state (**Fig. 3d**).

**Figure 3:**
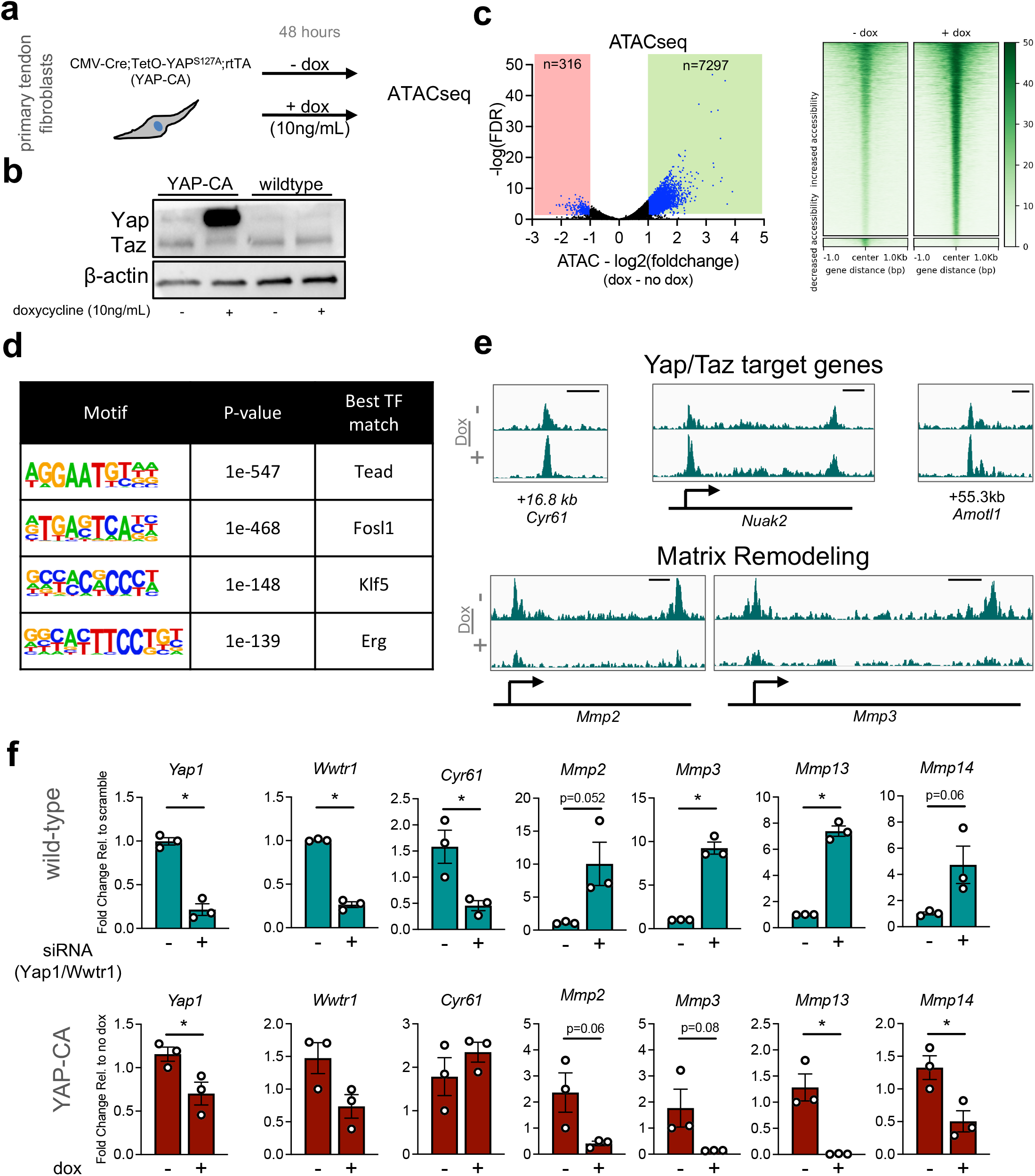
Yap overexpression maintains chromatin accessibility and represses matrix degradation pathways. (a) Schematic showing approach to examine the role of Yap in maintenance of chromatin accessibility. (b) Western blot showing robust overexpression of Yap following 48 hours of doxycycline treatment (10 ng/mL). (c) Volcano plot from ATACseq analysis showing differentially accessible genomic loci (adjusted p-value ≤ 0.05 and a fold-change of ≤ -1 or ≥ 1) in blue (n=3 biological replicates). Heatmaps of chromatin accessibility changes following blebbistatin treatment are also shown. (d) De novo transcription factor motif enrichment analysis from the genomic loci identified in (c). (e) Representative chromatin accessibility tracks showing an increase of accessibility of Yap/Taz target genes (*Cyr61, Nuak2, Amotl1*) and a reduction of accessibility of *Mmp2* and *Mmp3* following Yap overexpression. Genomic distance represents distance from gene TSS. Scale bar represents 1kb. (f) qRT-PCR analysis of tendon fibroblasts following loss- and gain-of-function of Yap/Taz (n=3 biological replicates). *p<0.05 evaluated by unpaired t-test. Error bars represent standard error.

Investigation of specific gene loci that changed in accessibility confirmed an increase of accessibility of Yap/Taz/Tead target genes (e.g., *Cyr61, Nuak2*, and *Amotl1*) (**Fig. 3e**). Interestingly, we also observed a marked reduction of accessibility for genes involved in matrix degradation (e.g., *Mmp2* and *Mmp3*) (**Fig. 3e**) demonstrating an epigenetic role of Yap/Taz in modulating matrix degradation pathways. To test whether Yap/Taz directly regulates expression of genes involved in matrix degradation, we administered siRNA targeting *Yap1* and *Wwtr1* (*Taz*) for 3 days and then analyzed gene expression by qRT-PCR. We found that depletion of Yap/Taz resulted in strong upregulation of *Mmp2, Mmp3, Mmp13*, and *Mmp14* (**Fig. 3f**). We further performed gain of Yap function assays using fibroblasts isolated from YAP-CA mice and administered doxycycline for 48 hours. Consistent with our siRNA data, overexpression of YAP repressed *Mmp2, Mmp3, Mmp13*, and *Mmp14* (**Fig. 3f**). Taken together, these data demonstrate that Yap/Taz are potent repressors of matrix degradation gene programs and that they act through direct epigenetic regulation of their respective genomic loci.

### Overexpression of YAP protects chromatin mechano-homeostasis following loss of cytoskeletal tension

Our data demonstrated that loss of cytoskeletal tension disrupted chromatin homeostasis (**Fig. 2c**), resulting in increased expression of matrix degradative gene programs (**Fig. 2b, f**), potentially through a loss of Yap/Taz signaling (**Fig. 2d**), and that Yap/Taz are potent regulators of expression of matrix degradative gene programs (**Fig. 3f**). This raised the interesting possibility that over-activation of Yap/Taz might serve as a counterbalance to the change in chromatin and transcriptional state that occurs with the loss of tensional homeostasis in cells. To test whether gain of Yap function could protect nuclear state following loss of tension, we overexpressed YAP in tendon fibroblasts from YAP-CA mice, administered blebbistatin for 6 hours, and then collected cells for RNA and ATAC analysis (**Fig. 4a**). Using qRT-PCR, we found that overexpression of YAP prevented the blebbistatin-induced downregulation of known Yap/Taz target genes (e.g., *Cyr61*) while also preventing the upregulation of matrix degradative gene programs (e.g., *Mmp2, Mmp3, Mmp10, Mmp13*, and *Mmp14*) (**Fig. 4b**). These data support that activating Yap signaling could protect the transcriptional state of the cell following de-tensioning. Interestingly, ATACseq analysis comparing Yap overexpressing cells with or without blebbistatin treatment revealed that non-muscle myosin inhibition still resulted in a strong reduction in chromatin accessibility even with Yap overexpression (**Fig. 4c, d**). While these data are consistent with our prior blebbistatin ATACseq data from wildtype fibroblasts (**Fig. 2c**), we intriguingly no longer found Tead as the leading candidate in the motif enrichment (**Fig. 4e**), indicating that alternative Yap/Taz independent pathways could be the contributing to the loss of chromatin accessibility under these conditions.

**Figure 4:**
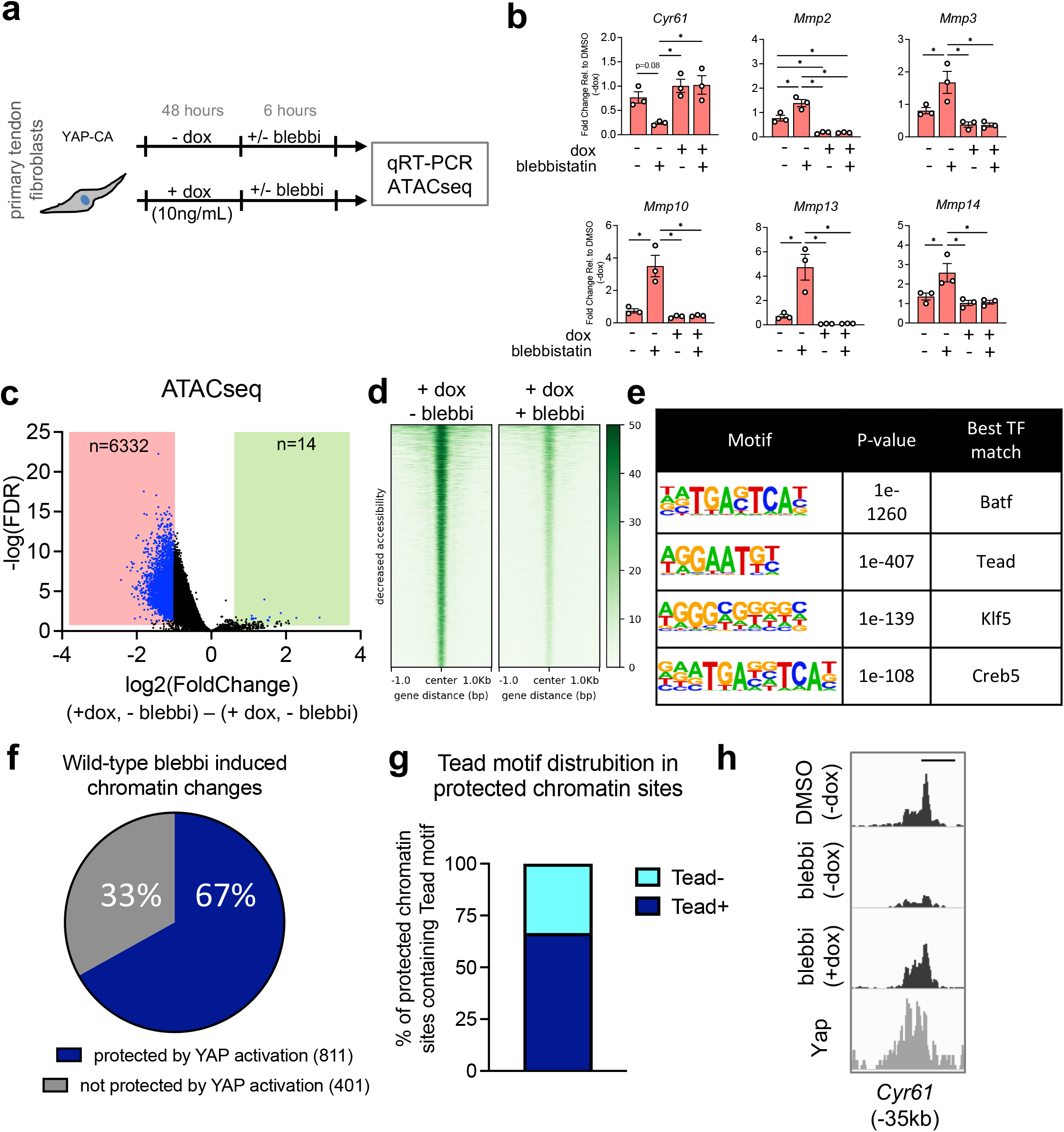
Activation of Yap signaling protects against catabolic changes from loss of cellular tension by maintaining chromatin accessibility at mechano-responsive loci. (a) Approach to examine whether Yap gain-of-function could attenuate transcriptional and epigenetic changes following loss of tension. (b) qRT-PCR analysis of tendon fibroblasts following Yap overexpression and blebbistatin treatment (n=3 biological replicates). (c) Volcano plot from ATACseq analysis showing differentially accessible genomic loci (adjusted p-value ≤ 0.05 and a fold-change of ≤ -1 or ≥ 1) in blue (n=3 biological replicates). (d) Heatmaps of chromatin accessibility changes following blebbistatin treatment. (e) De novo transcription factor motif enrichment analysis from the genomic loci identified in (c). (f) % of chromatin accessibility changes in wild-type tendon fibroblasts following blebbistatin treatment that no longer changed in accessibility following Yap overexpression. (g) distribution of the Tead motif in the protected chromatin sites identified in (f). (h) representative ATAC data showing protection of chromatin accessibility changes in Yap overexpressing cells following blebbistatin treatment. Genomic distance represents distance from gene TSS. Scale bar represents 1kb. *p<0.05 evaluated by unpaired t-test. Error bars represent standard error.

To assess whether Yap overexpression in the context of blebbistatin treatment protected the accessibility of Tead chromatin binding sites, which were inhibited in wild type cells (**Fig. 2c**), we specifically examined the accessibility profiles of those chromatin loci in Yap overexpressing cells treated with blebbistatin compared with cells only overexpressing Yap. Interestingly, we found that 67% (n=811) of the chromatin loci that changed accessibility in the wild-type cells with blebbistatin no longer changed chromatin accessibility in the Yap overexpressing cells following blebbistatin treatment (**Fig. 4f**). This demonstrates that activation of Yap protects the chromatin state following cellular de-tensioning. We next examined these 811 chromatin loci for presence of the Tead DNA binding motif and found motif enrichment in 2/3rds (66.5%) of the chromatin loci (**Fig. 4g**). Finally, we cross-examined the protected ATAC loci with publicly available Yap ChIPseq data and found strong overlap of Yap occupancy at the sites of protected chromatin loci following loss of tension (**Fig. 4h**). Collectively, these data reveal that overexpression of Yap protects both chromatin accessibility and transcriptional state following loss of tensile cues in fibroblasts.

## Discussion

This study reveals, for the first time, that tensile cues actively maintain chromatin mechano-homeostasis through a Yap and Taz axis. Upon disruption of tensional homeostasis *in vivo*, nuclei round up and become disorganized, and matrix degradation pathways are activated. Similarly, with a loss of cell tension *in vitro*, the mechano-responsive chromatin state changes, resulting in the activation of matrix degradation pathways. Using paired ATAC/RNAseq we identify Yap/Taz/Tead as central epigenetic and transcriptional regulators of the nuclear response to loss of tension. Depletion of Yap and Taz elevates gene expression of MMPs, similar to that which occurs with cytoskeletal de-tensioning, while overexpression of Yap represses MMP gene expression, even when the cytoskeleton is de-tensioned. These exciting results demonstrate a direct epigenetic role of Yap and Taz proteins in the epigenetic repression of genes involved in matrix degradation. Finally, we show that overexpression of Yap preserves chromatin accessibility, and their respective expression, at distinct mechano-responsive genomic loci following cellular de-tensioning.

Disruption of tensional homeostasis is an important factor in the progression of a wide variety of chronic diseases, including cancer, cardiovascular disease, and tendinopathies (27– 29). Our finding that inhibition of tensional homeostasis activates matrix degradation through an epigenetic and transcriptional Yap and Taz axis provides a novel mechanism by which disruption of tensile cues leads directly to persistent matrix degradation and/or remodeling. Interestingly, we find that loss of tensile cues reduced chromatin accessibility, which is consistent with recent work showing that reducing matrix stiffness, a different mechanical input to the cell, also reduced chromatin accessibility (20, 21). While these previous studies did not primarily focus on Yap/Taz, a distinct Yap/Taz signature was observed in their datasets, suggesting that Yap/Taz could be principal mechano-epigenetic regulators of chromatin state. Furthermore, consistent with our observations, previous work demonstrated that activation of signaling pathways which block Yap/Taz signaling (e.g. cAMP) initiate matrix degradation (30, 31). These studies, taken together with our findings, demonstrate that the nucleus is exquisitely sensitive to mechanical cues, with increased mechanosignaling (i.e., tension, matrix rigidity) promoting global chromatin accessibility. However, additional studies are needed to determine whether the changes in chromatin state are due to transcription factor abundance in the nucleus (i.e., Yap/Taz) or to direct mechanical control of chromatin accessibility and transcription via nuclear (and so chromatin) deformation (13). Such insights into these mechanisms may very well inform critical epigenetic programs that regulate disease progression.

Tendinopathy, particularly from overuse, is thought to arise from tissue microdamage. This microdamage, in turn, impairs transmission of tensile mechanical signals through the tissue, resulting in under-loading of endogenous tendon cells (7, 32). The disease often leads to aberrant cell and matrix responses, such as chondroid metaplasia of the tendon, further attenuating ECM-to-cell strain transfer (8, 33). Interestingly, tendinopathic specimens also contain myofibroblasts and appear to undergo chronic remodeling, which may be an attempt by the resident tendon fibroblasts to re-tension their matrix to re-establish tensional homeostasis(27). Ongoing failure to restore the proper tensional set point is likely a contributing factor in the progression of the disease. Our finding that acute de-tensioning results in a marked induction of a catabolic transcriptional program and that the Yap/Taz/Tead axis protects chromatin state following loss of tension, preventing this catabolic response, could inform mechanisms that regulate the initiation and early progression of tendinopathy. Beyond the catabolic response, acute de-tensioning also elicits cell death which can be protected through inhibition of cell contractility (4, 5). Therefore, it is likely that control of cellular tension is required to maintain cell fate and tissue homeostasis. Importantly, our findings offer new insight towards the development of novel mechano-therapeutic targets that could directly regulate this balance in cellular tension to attenuate disease progression.

While we found that both cellular de-tensioning and Yap/Taz regulate expression of MMPs, we only found changes in chromatin accessibility at MMP genomic loci when overexpressing a mutated and constitutively-active Yap (**Fig. 3e**). While the relationship between ATAC signal and gene expression is thought to be linear (i.e., increase in accessibility is associated with increased expression) (34), temporal analysis of these assays during dynamic changes, such as single factor perturbations, is less clear (35). We performed ATAC/RNAseq at 6 hours after blebbistatin and Y27632 treatment, and so it is possible this short duration was too early for observable and measurable ATAC differences to occur. Alternatively, it is possible that additional transcriptional machinery downstream of Yap/Taz could also be regulating MMP expression, further convoluting the mechanism by which cellular tension regulates MMP expression. Delineating the precise contribution of ATAC signal and MMP gene expression will require future temporal analysis of these genomic loci in response to prolonged and altered mechanical signaling.

Collectively, our findings provide novel transcriptional insights into the mechanisms by which applied tension leads to mechano-signaling that directs cell behavior on an epigenetic basis. This epigenetic program, particularly chromatin accessibility and suppression of matrix catabolism, is mediated by Yap/Taz. These findings open up new avenues for further investigation of these understudied biological processes in the context of tissue formation, homeostasis, and disease progression.

## Methods

### Animals

All animal procedures were approved by the University of Pennsylvania’s Institutional Animal Care and Use Committee. The following mice were used: Tg(CMV-cre)1Cgn/J (Jackson Laboratory, strain #006054), TetO-Yap^S127A^ (gift from Carmago and Calloway labs), and Gt(ROSA)26Sor^tm1(rtTA,EGFP)Nagy/J^ (Jackson Laboratory, strain #005670), and wild-type CD-1 IGS (Charles-River, strain #022).

### FDL resection surgeries

The FDL tendon consists of a proximal region that attaches to the flexor digitorum longus muscle, a wrap-around region that glides along the distal-medial edge of the tibia and is surrounded by a synovial sheath, and the distal region in the forefoot that attaches to the phalanges of digits 2-5. In the FDL resection procedure, first mice (2-3 months old) were anesthetized with isoflurane (1–3%), given pre-surgical analgesia, and sterilely prepped. The distal FDL was exposed and a 3 mm segment proximal to the bifurcation point was resected in order to de-tension the proximal region of the tendon. The skin was closed with 4-0 nylon suture and the animal was returned to unrestricted cage activity. This model was used to induce acute de-tensioning on the tendon while also assessing the proximal region that is less effected by inflammation and granulation tissue of the healing tissue at the resected site in the forefoot.

### Tissue sectioning and RNA *in situ* hybridization

Following euthanasia, hindlimbs were harvested, fixed in RNase-free 4% paraformaldehyde (PFA) for 24 hours with the ankle at 90 degrees of flexion, decalcified in ethylenediaminetetraacetic acid (EDTA) containing 2% PFA for 7 days, and embedded in optimal cutting temperature (OCT) compound. Tape-stabilized (Cryofilm 2C, Section-Lab Co. Ltd.) sagittal sections were made and then stained for Mmp3 (Advanced Cell Diagnostics, Cat No. 480961) using the RNAScope 2.5 HD Assay (Advanced Cell Diagnostics, Inc.) using the custom pretreatment reagent (Cat No. 300040) according to the manufacturer’s protocol.

### Primary tendon fibroblast isolation and cell culture

Tail tendons were extracted from both 6-8 week old male and female CD1 wildtype mice. Tendons were digested with an enzymatic solution (DMEM, 0.2 mg/mL Liberase DL, and 100 U/mL DNase I, Thermo-Fisher Scientific) for 60 minutes at 37°C. Digestive solution was inactivated with DMEM containing 10% FBS and antibiotics-antimycotics. Cell and tissue suspensions were passed through a 70 μM filter and centrifuged at 300g at 4°C for 10 minutes. Cells were then resuspended in cell culture medium (DMEM supplemented with 10% FBS and antibiotics-antimycotics) and then seeded on collagen I coated tissue-culture plastic. Cells were used between passage 1-3. Blebbistatin (Caymen Chemicals) was used at 10 uM in DMEM supplemented with 0.1% FBS with antibiotics-antimycotics. Y27632 (Caymen Chemicals) was used at 30 uM in DMEM supplemented with 0.1% FBS with antibiotics-antimycotics.

### RNAi knockdown

RNAi was carried out with SMARTpool: ON-TARGETplus siRNA’s specific for mouse *Yap1* and *Wwtr1* (Dharmacon) or scramble non-targeting siRNA as a control. Primary mouse tendon fibroblasts were transfected with Lipofectamine RNAiMAX (Thermo Fisher Scientific) in DMEM supplemented with 10% FBS and antibiotics-antimycotics for 72 hours, after which they were reseeded on tissue-culture plastic at lower density (7,500 cells/cm^2^).

### Protein lysis and western blotting

For protein analysis, cells were lysed with RIPA buffer (Thermo Fisher Scientific) supplemented with Halt protease and phosphatase inhibitor (Thermo Fisher Scientific). Protein concentration was quantified with the Pierce BCA Protein Assay kit (Thermo Fisher Scientific). A total of 5 ug of protein was loaded per sample. Samples were loaded onto 4-15% gradient SDS-PAGE gels, then transferred to 0.2 um pore-size PVDF membranes (Bio-Rad). Blots were blocked in blocking buffer (5% milk, 0.1% Tween-20 in 1x TBS) and then treated with primary antibodies (YAP/TAZ, anti-rabbit, Cell Signaling, 8418 and beta-Actin, anti-mouse, Cell Signaling, 3700) overnight at 4°C. Next, blots were washed 3x in 1x TBS with 0.1% Tween-20. Blots were then incubated in species specific secondary HRP-conjugated antibody (anti-rabbit, Promega, W4011 and anti-mouse, Promega, W4021) for 1 hour at room temperature. Blots were then washed 3x in 1x TBS with 0.1% Tween-20. Protein bands were visualized using Super Signal West Pico Plus (Thermo Fisher Scientific), and images were acquired using a ChemiDoc XRS Imaging System (BioRad).

### RNA isolation, cDNA synthesis, and qRT-PCR

mRNA isolation was carried out using the Qiagen mRNeasy Plus Mini Kit (Qiagen). RNA concentration was quantified using a Nanodrop spectrophotometer. cDNA was synthesized using the SuperScript VILO IV kit (Thermo Fisher Scientific). Quantitative-real time PCR (qRT-PCR) was completed using Fast SYBR reagents (Thermo Fisher Scientific) and analyzed using a QuantStudio 6 Pro (Applied Biosystems). Primer sequences are provided in Table S1.

### RNA sequencing and analysis

RNA quality was evaluated using a BioAnalyzer. RNA libraries were prepared using Illumina truSeq polyA high-throughput kit and single-end reads were sequenced to 100 base pairs using an Illumina NovaSeq. Reads were aligned to the mm10 genome using Hisat2 (36). Picard was used to generate bam files and sorted by chromosomal coordinates. FeatureCounts was used to generate count files (37). Deseq2 was used for differential expression analysis between groups (n=3 biological replicates/group) (38). Pathway analysis was completed using Panther.

### ChIPseq analysis

Publicly available Yap ChIPseq data was downloaded from GSE83863 (24). Sequenced reads were aligned to the mm10 build of the mouse genome using Bowtie2 with default settings (39). SAM files were converted to BAM and were sorted by chromosomal coordinates using Picard SortSam. Duplicates were removed using Picard Mark Duplicates. BAM files were then used to call peaks using MACS2, with default settings with a q threshold of 0.05 (40). To generate BigWig files, deepTools BamCoverage was used with default settings and a bin size of 10 bp (41). Integrative Genome Viewer was used to visualize BigWig files on the mm10 genome track.

### ATACseq and analysis

50,000 cells were subjected to Assay for Transposase Accessible sequencing (ATACseq) following the published protocol (42). DNA fragment size was analyzed using a Fragment Analyzer. Libraries were sequenced to 50 base pairs from both ends using an Illumina NovaSeq. Raw reads were aligned to the mm10 genome using Bowtie2. Picard and Samtools were used to generate bam files and to filter out duplicates and mitochondrial reads. Peaks were called using MACS2 using the following options: ““--keep-dup all -q 0.01 -no model”.

Diffbind was used to identify differentially accessibility chromatin loci between groups (n=3 biological replicates/group) (42). Motif analysis was completed using the findMotifsGenome.pl command within Homer (43).

## Acknowledgements

We would like to acknowledge the following funding sources: National Institutes of Health (R00AR067283, R01AR075418, and P30 AR069619) and the Department of Veterans Affairs (IK6 RX003416). Additional support was provided by a Gilliam Fellowship (GT13516) from the Howard Hughes Medical Institute. We would like to thank the Penn Genomic Sequencing Core for their assistance with RNA/ATACseq and the Penn Microscopy Core. We would also like to thank Drs. Fernando Carmago and Jenna Galloway for supplying the YAP-CA mice. We would finally like to thank all members of the Mauck and Dyment labs for thoughtful discussion and feedback during the development of this project.

## Author Contributions

All authors contributed to conceptualization and experimental design. DLJ, RD, XJ, MKE, EB, and MN performed the experiments. DLJ, RD, XJ, EB, RLM, and NAD analyzed and interpreted data. DLJ, RLM, and NAD drafted the manuscript. All authors contributed to editing the manuscript.

## Data Availability

Raw and analyzed ATACseq and RNAseq data generated in this study are available through the Gene Expression Omnibus under the GEO accession GSE207896.

